# Distinct mobility patterns of BRCA2 molecules at DNA damage sites

**DOI:** 10.1101/2023.02.20.527475

**Authors:** Maarten W. Paul, Jesse Aaron, Eric Wait, Romano M. Van Genderen, Arti Tyagi, Hélène Kabbech, Ihor Smal, Teng-Leong Chew, Roland Kanaar, Claire Wyman

## Abstract

BRCA2 is an essential tumor suppressor protein involved in promoting faithful repair of DNA lesions. The activity of BRCA2 needs to be tuned precisely to be active when and where it is needed. Here, we quantified the spatio-temporal dynamics of BRCA2 in living cells using aberration-corrected multifocal microscopy (acMFM). Using multicolor imaging to identify DNA damage sites, we were able to quantify its dynamic motion patterns in the nucleus and at DNA damage sites. While a large fraction of BRCA2 molecules localized near DNA damage sites appear immobile, an additional fraction of molecules exhibits subdiffusive motion, providing a potential mechanism to retain an increased number of molecules at DNA lesions. Super-resolution microscopy revealed inhomogeneous localization of BRCA2 relative to other DNA repair factors at sites of DNA damage. This suggests the presence of multiple nanoscale compartments in the chromatin surrounding the DNA lesion, which could play an important role in the contribution of BRCA2 to the regulation of the repair process.

## INTRODUCTION

The activity of proteins needs to be controlled spatially and temporally. This is especially true for proteins that maintain the integrity of the genome through DNA repair reactions. These proteins must act at the right time and place to repair DNA damage. They need to cooperate or compete with many other proteins at the same genomic location, such as proteins involved in replication and transcription. The critical parameters that define the activity of these proteins in cells are their concentration and their diffusion rate. Simultaneously, both transient, non-specific and specific interactions determine their spatial organization. These specific and non-specific interactions can be with other proteins but also with other biomolecules such as DNA and RNA.

BRCA2 is an essential DNA repair protein that is present in a low concentration in the cell nucleus. It is found in a complex with several other DNA repair proteins, such as PALB2 and RAD51, which are all required at sites of DNA damage lesions repaired by homologous recombination (1). The central function of BRCA2 is to exchange the single-strand DNA binding protein complex RPA with RAD51 on the resected DNA ends of DNA double-strand breaks (DSB). Additionally, the resolution of stalled replication forks and of interstrand-crosslinks by homologous recombination requires localized activity of BRCA2 at damage sites (2, 3). As an important tumor suppressor, BRCA2 is an extensively studied protein, where BRCA2 mutations are associated with hereditary breast and ovarian cancer (4). However, the biophysical properties of BRCA2 that drive its function in cells are not fully understood. Here, our objective is to define the dynamic behavior of BRCA2 around DNA lesions to understand the critical parameters that are required for BRCA2 to act properly.

DNA-associated proteins can exhibit different modes of diffusion that contribute to their different functions (5, 6). For example, recently it has been reported that the transcriptional regulator CTCF maintains transient binding zones within the cell nucleus, which could be a mechanism to improve its target search efficiency in the nuclear volume (7). Although many nuclear proteins, such as transcription factors, are involved in direct and specific interactions with DNA (sequence motifs, specific structures), it appears that BRCA2 does not depend on this type of interaction for its immobilization at damage sites, but rather on (transient) protein-protein interactions (8). The immobilization and accumulation of BRCA2 is essential for its function, but what interactions are relevant for this behavior and how this correlates with its function is largely unknown. Several reports indicate a crucial interaction between BRCA2 and PALB2, and indeed BRCA2 accumulation at damage sites depends on the interaction with PALB2 (9), whereas localization of PALB2 itself is influenced by BRCA1 (10) and the ubiquitin ligase RNF168 (11). Recent data suggests that PALB2 chromatin interaction depends on chromatin associated proteins (12, 13), hence those chromatin interactions could also contribute to the diffusive patterns that have been observed for its binding partner BRCA2 (8, 14).

BRCA2 accumulates at DNA damage into so-called foci, like several other homologous recombination repair proteins. It is not yet fully understood how redistribution occurs and how BRCA2 proteins organize themselves at damage sites. In this study, we applied 3D single-molecule multiplane imaging to define the quantitative behavior of endogenous BRCA2 at fluorescently marked DNA damage sites. This revealed that at DNA damage sites, immobilization of BRCA2 increases, but also that those molecules that diffuse near DNA lesions do so with increased confined mobility. Quantification of the number of BRCA2 molecules in the nucleus and at damage sites indicated that only tens of BRCA2 molecules are present per focus, while super-resolution microscopy confirmed the localization of BRCA2 in nanoscale clusters within foci.

## MATERIAL AND METHODS

### Cell culture

Mouse ES cells were maintained on 0.1% gelatin (Sigma) coated plates in mouse ES cell medium (DMEM, 40% BRL-conditioned medium, 10% FCS (Capricorn Scientific), supplemented with pen-strep, non-essential amino acids (Lonza) and leukaemia inhibitory factor (1000 U/ml) and 0.1 mM β-mercaptoethanol). For the ac-MFM experiments cells were maintained for several passages prior to the experiments in DMEM Knockout medium (Thermofisher) supplemented with 10% FCS (Sigma, F2442), pen-strep, LIF, NEAA and 0.1 mM β-mercaptoethanol. mES cells expressing BRCA2-HaloTag previously generated and described in (1). Mouse ES cells expressing BRCA2-EGFP were previously generated and described in (2).

### DNA constructs

The EGFP-tr53BP1 construct was assembled using Gibson Assembly with a fragment of human 53BP1 (amino acids 1221-1709, Addgene #69531) (4) that was cloned in a EGFP Piggybac vector (3).

### Generation of cell lines

The EGFP-53BP1trunc Piggybac vector was transfected in BRCA2-HaloTag cells together with the CMV-Hypbase Piggybac transpose plasmid using Lipofectamine 3000 according to the manufacturer’s protocol. Cells were selected with puromycin for 1 week after which the selected population was used for the experiments.

### Sample preparation of live cell imaging

Between 20,000 and 40,000 cells were seeded the day before the imaging experiment on 8-well ibidi dishes coated with 25 μg/ml laminin (Roche, 11243217001). When indicated, cells were treated with 1 μg/ml mitomycin C (Sigma, M0503). For HaloTag labeling, the medium was replaced and, subsequently, the cells were labeled with 50 nM JF549-HaloTag dye, unless a different concentration is indicated, followed by washing with PBS and replacement with fresh medium. Imaging was done in FluoroBrite DMEM medium (ThermoFisher, A1896701) complemented with 10% FCS, pen-strep, LIF, NEAA, 0.1 mM β-mercapto-ethanol.

For FRAP experiments cells were labelled with 250nM of JF549-HaloTag ligand and cells were labelled and washed as described above. For 2D tracking experiments in wild-type and 53bp1 knockout cells, cells were labeled with 0.5 nM JFX549 for 10 minutes.

### Sample preparation for immunofluorescence

The cells were grown in 8-well glass bottom dishes (Ibidi) or 24 mm round coverslips (#1.5H, Marienfeld, 0117640), coated with laminin as described above. For fixation cells were washed with PBS once and fixed with 4% PFA (Thermo-Fisher) in PBS for 10 minutes. Cells were washed with PBS with 0.1% triton for 3 times followed by two 10-minute washing steps. Cells were blocked with 3% BSA in PBS for 20 minutes followed by incubation of the primary antibodies in blocking buffer. After washing with PBS with 0.1% Triton, cells were incubated with fluorescently labeled secondary antibodies for 1 hour at room temperature.

For immunostaining of 53BP1 a rabbit polyclonal antibody was used (1:1000, Novus biologicals, NB100-304) together with anti-rabbit F(ab′)2 fragment conjugated with CF568 (1:1000, Sigma, SAB4600310).

For dSTORM imaging of HaloTag protein fusions, cells were incubated with 500 nM AF647-HaloTag ligand (kind gift from Luke Lavis) together with the primary antibodies in blocking buffer for 2 hours at room temperature.

For STED imaging, mES cells expressing BRCA2-EGFP (2) were treated with Mitomycin C (1 μg/ml) for 2 hours. Subsequently, the cells were fixed 2 hours later with 4 % PFA. Cells were immunostained as described above using anti-GFP nanobody conjugated with STAR635P (1:500, FluoTag X4, 0304-Ab635P, NanoTag). 53BP1 was visualized with anti-53BP1 antibody (1:2000, Novus Biologicals, NB100-304) and secondary anti-rabbit Alexa594 (1:2000, ThermoFisher). Samples were mounted in ProLong Gold mounting medium on object glasses.

### Microscope setup and image acquisition

*Ac-MFM setup* The ac-MFM system was as described by Abrahamson et al (5). In summary, specimens were imaged on a Nikon TiE epifluorescence microscope with a 100x 1.45NA Plan Apo objective lens (Nikon) and illuminated by 3-6kW/cm^2^ 561 nm laser excitation. Fluorescence was collected and passed through a custom multifocal grating and color correction grating as described by Abrahamsson et al. Nine focal planes were imaged onto two EMCCD detectors (iXon DU897, Andor) with a frame rate up to 30 Hz. The focal position of the microscope was maintained by active stabilization.

Every day, prior to the imaging experiments, a sample with mounted 100 nm Tetraspeck beads (ThermoFisher) was used to correct for chromatic aberrations, to measure the z-slice separation distance, to determine relative z-plane detection efficiency, and to align image channels. A Tokai Hit microscope stage holder was used to maintain cells at 37C with 5% CO_2_. Movies of 2000 frames were recorded, where in every 50^th^ frame the GFP signal was also recorded on the second camera detector. The sample was continuously illuminated with 561nm light and constant low 405 nm light for photoactivation of Halo-JF549.

*Confocal imaging and FRAP* Experiments were conducted on a Zeiss Elyra PS1 system with a Tokai Hit stage incubator calibrated at 37 °C and 5 % CO_2_ using a 63x objective (NA 1.4 Plan Apo). For both FRAP experiments (**Figure 1**) and quantitative confocal experiments (**Figure 3**) a GaAsP detector was used. FRAP experiments were acquired using a square ROI of 1.14x1.14 µm, which was scanned at a 0.2 s interval, with 20 frames before bleaching, 4 frames bleaching at full laser power, followed by 230 frames to record recovery. Cells that moved too much during acquisition were excluded from the analysis. Mean intensity from the bleach ROI was measured over time. The intensity traces were normalized by the mean intensity of frame 10-20 before bleaching. The data in **Figure 1A** shows the average normalized intensity trace of multiple experiments, whereas the shading represents the standard error of the mean intensity.

**Figure 1:**
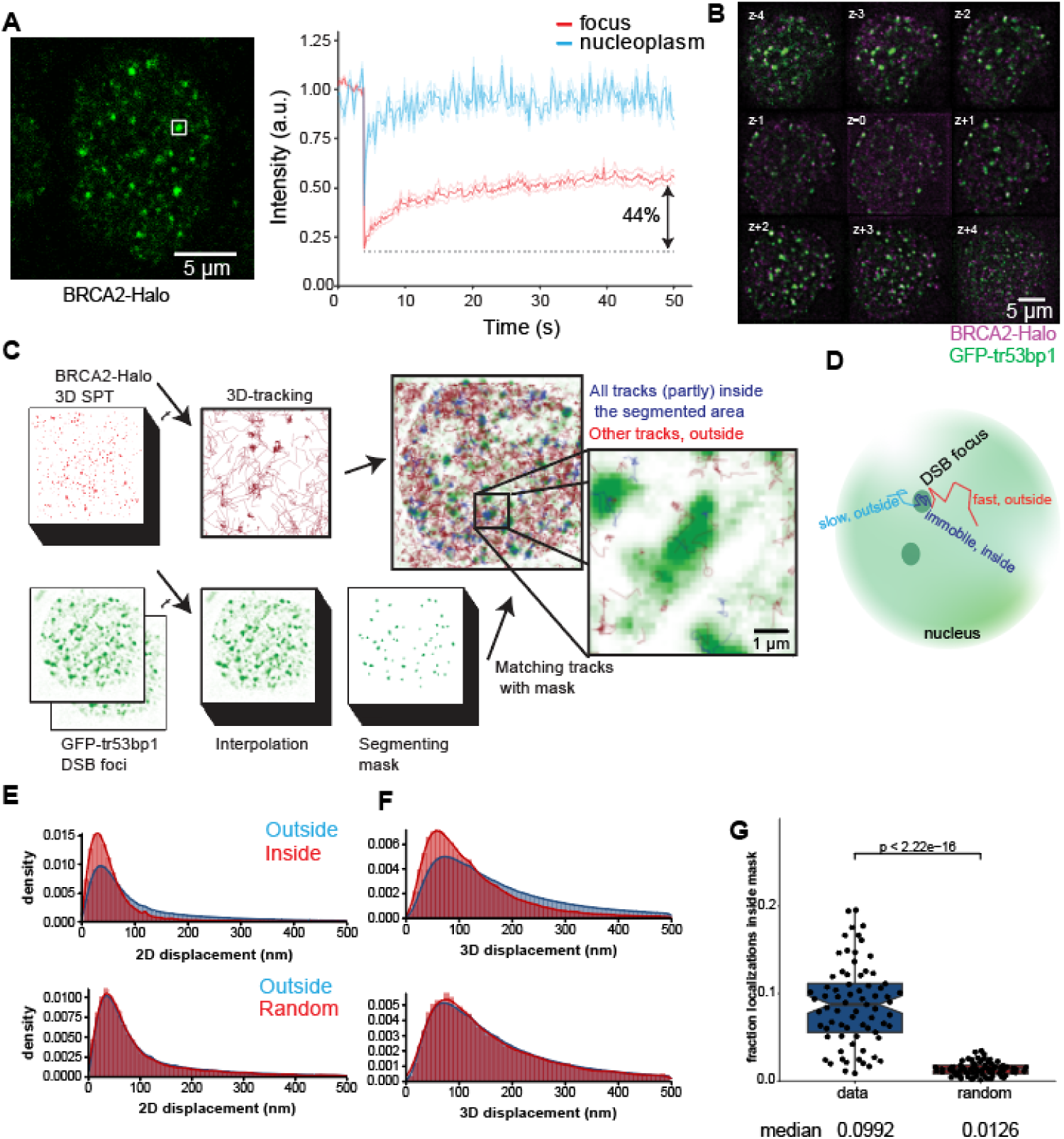
Analysis of the mobility of BRCA2 at DNA damage sites. **A** Spot-FRAP of BRCA2-HaloTag cells treated with mitomycin C by photobleaching a focus or region in the cytoplasm (3 independent replicates, focus n=40, nucleoplasm n=20 cells, shading around the curve indicates SEM) **B** Example of a single multi-focal plane image showing 9 z-plane projections spaced 420 nm apart; the signal of BRCA2-Halo (magenta) and EGFP-tr53BP1 (green) is shown. **C** Approach for analysis of molecule tracks in and outside the foci, by matching the localized molecules to the binary mask of the damage sites defined by the 53BP1 signal. **D** All track segments in the data set are annotated as being in or outside the DSB foci and having a fast, slow, or immobile state. **E,F** Distribution of mean x-y (2D) and x-y-z (3D) displacements for track segments outside versus inside the DSB foci mask. The plots below show the distribution but for a random mask. **G** Fraction of localizations in and outside the defined mask for the data compared to the random mask. The data consist of 4 independent experiments of 73 cells in total. See Movies S1-3 for movies of this data. p-values are calculated from an unpaired 2-sample Wilcox rank sum test.

*dSTORM* Imaging was performed using a Zeiss Elyra PS1 system using a 100x 1.46NA Korr α Plan Apochromat objective. 561 and 642 100mW diode lasers were used to excite the fluorophores together with respectively a BP 570–650 + LP 750 or LP 655 excitation filter. dSTORM imaging was performed using near-TIRF (HiLo) settings, while the images were recorded on an Andor iXon DU 897, 512x512 pixel EMCCD camera.

For drift correction and channel alignment 100 nm Tetraspeck beads (ThermoFisher) were added to the sample. To perform dSTORM imaging, an imaging buffer was prepared containing 40 mM MEA (Sigma), 0.5 mg / ml glucose oxide (Sigma), 40 μg/ml Catalase (Sigma) and 10% w / v glucose in Tris pH 8.0. dSTORM images were acquired sequentially, starting with Alexa 647 staining followed by CF568 imaging. At least 10,000 images were acquired at an interval of 33ms per frame.

The raw data was analysed by ZEN software. The molecules were localized, and a drift correction was applied using image correlation in the two images separately. Localizations that were present within 20 nm of each other in the 5 subsequent frames were grouped. Subsequently, the two images were combined, and the channels were aligned laterally using 100 nm Tetraspeck beads that are deposited into the sample.

For blinking correction of the plots in **Figure S8** we made use the pairwise distance distribution correction algorithm (DCC) (6). ROIs with ungrouped localizations were loaded into Matlab and processed with the DCC script taking into account the photon counts as obtained from the ZEN software.

*STED* The samples were imaged on a Leica SP8 tauSTED. Star635 signal was excited with a white-light laser tuned at 632 nm and 775 nm depletion laser with a 2D STED depletion pattern, whereas Alexa594 was imaged with the same depletion laser, but excited with the white light laser tuned at 561 nm. STED images were deconvolved using Huygens software (SVI).

### Image analysis for ac-MFM

Raw images from the ac-MFM microscope were converted to 3D stacks and corrected for chromatic aberrations in MATLAB (https://github.com/aicjanelia/MFM) and processed using background subtraction and Richardson-Lucy deconvolution (7px rolling ball, 5 iterations, respectively). Subsequently, molecules were localized and tracked in 3D using the TrackMate plugin (7) in ImageJ after pre-processing using and Gaussian blur denoising (0.7px radius). The molecules were linked at a maximum frame-to-frame distance of 500 nm.

EGFP-tr53bp1 movies, that were used to define the DNA damages sites, were processed in Matlab, where frames were identified that contained EGFP signal, the signal in intermediate frames was interpolated linearly based on the image frame before and after. The foci signal was identified using Laplacian of Gaussian edge detection and a binary mask was created.

Next, for every point of the tracks it was determined whether it was in or outside the mask. To compare the results to a random situation, data sets were generated by randomly redistributing the identified 3D objects at the first frame of the movie, along the x-y axes in the image plane within the convex hull surrounding the centre of mass of all foci. The MATLAB routines used for the analysis can be found in https://github.com/maartenpaul/MFManalysis_Matlab/ (based on original code repositories: https://github.com/aicjanelia/MFM and https://gitlab.com/aicjanelia/visitor-maarten-paul). Subsequently a script in R was used for processing of the tracks and plotting of the data (https://github.com/maartenpaul/MSDanalysis_MFM). Python code to segment the tracks into different mobile states using deep learning and to estimate the mean square displacement and moment scaling spectrum of the individual tracklets (8) was incorporated into Rstudio using the reticulate package (9).

For further analysis, track(let)s presenting at least one frame of their lifetime inside the mask were considered associated with the DNA damage sites.

The angle (𝜃𝜃) between two displacements within the track(let)s was calculated as described in Hansen (2019) (7). For the angle analysis displacements, immobile tracklets and x-y displacements below 100 nm were excluded from the analysis.

For a single track(let) with 𝑟(𝑡) = (𝑥(𝑡), 𝑦(𝑡)) its coordinates and 𝑇 ≥ 8 its total track length, the mean squared displacement (MSD), at different time lags 𝛥𝑡, was computed as follow:

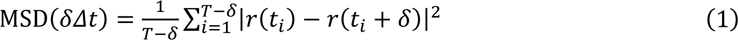

The apparent diffusion constant *D_app_* and anomalous exponent 𝛼𝛼 were estimated by fitting a linear regression on the logarithm form of the MSD power-law equation (10) (using the first 4 MSD fitting points):

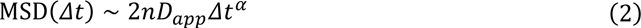

where 𝑛 is the number of dimensions.

The radius of confinement 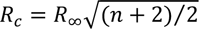 was measured after estimating the plateau 𝑅_∞_ of the MSD curve (11). The latter was estimated by fitting the following equation, using the first 20 MSD fitting points and the “curve_fit” function from the Python SciPy library:

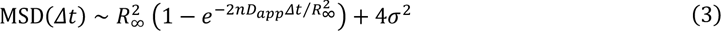

with 𝜎^2^ the experimental noise level.

### Image analysis dSTORM

For quantification of BRCA2 clusters, 53BP1 foci were identified by applying an Otsu threshold on 53BP1 images and saved as ImageJ ROIs. The ROIs were imported in R and BRCA2 localizations that are within the ROIs were clustered using the RSMLM package using the ToMaTo clustering algorithm (r=50, threshold=0.1) (13). To remove spare localizations clusters with less than 5 localizations were filtered out. Shape features describing these clusters were determined using the SMoLR package (14). The size of clusters is defined as the mean of the FWHM of the major and the minor axis of the localizations within the cluster. The distance between the clusters is determined as the distance between the centre of mass of the two clusters within the same segmented 53BP1 focus.

### Quantification of cellular concentration of BRCA2

HaloTag-GST (61 kDa) (Promega, G4491) was labelled with JF646 for 4 hours at 4 ° C on ice with 3x excess HaloTag ligand (buffer: 1x PBS, 20% glycerol, 1 mM DTT). Followed by removal of the unreacted HaloTag ligand using a Zeba desalting column (7K MWCO, 0.5 ml, ThermoFisher, 89882). Using SDS-PAGE it was confirmed that the labelled protein was recovered after labelling and elution from the column. The concentration of the labelled protein was validated using Nanodrop and on SDS-Page using a BSA with known concentration. Subsequently this labelled protein was used as reference to estimate the amount of BRCA2 per cell.

The nuclei of live cells were segmented in 3D using a routine in TrackMate (12), which uses Stardist (15) for 2D segmentation of nuclei, followed by connecting the matching nucleus segmentation in the 3D image stack. A 3D label image was exported and used in a custom-made routine developed in CellProfiler (16). The label image was used to generate 3D nuclei objects and was used to determine the nuclear volume and integrated intensity of the BRCA2-halotag intensity. The BRCA2 foci were segmented using an adaptive threshold in 3D and volume and intensity were measured. The CellProfiler pipeline and accompanying R code for processing and plotting the data is available at https://github.com/maartenpaul/Halo_cell_quant .

The integrated intensity (a.u.) was converted to concentration and number of molecules by comparing the intensity to a titration curve of the labelled HaloTag-GST::JF646 standard protein. Using this titration curve the intensity per pixel can be converted to the local concentration of HaloTag protein. Assuming that the segmented nuclei are much larger than the confocal volume per pixel this conversion can be used to estimate the concentration of BRCA2 in the nuclei. Subsequently, the concentration can be converted to number of molecules per nucleus by dividing the volume and Avogadro’s number.

For in-gel analysis of in vivo labelled BRCA2-HaloTag, HaloTag was labelled with JF646-HaloTag ligand in live cells grown on plates. Subsequently cells were trypsinized and the cell concentration was determined. Cells were resuspended at 25,000 cell/µl and treated with benzonase in buffer with protease inhibitors (2 mM MgCl2, 20 mM Tris pH 8, 10% glycerol, 1% triton X-100 and 25 U/ml benzonase) and subsequently lysed by adding 2x Laemmli buffer without bromophenol blue, at equal volume. Cell lysates were run on 3-8% SDS-Page Tris-Acetate gel. The fluorescently labelled proteins were imaged in the gel using a Typhoon gel imager (Amersham).

## RESULTS

Previous studies have shown, that BRCA2 appears more immobile after induction of DNA damage (8, 14). However, those studies did not investigate the spatial distribution of immobilized BRCA2 after induction of DNA damage. BRCA2 is involved in the repair of different types of lesions that also involve RAD51-mediated homology recognition and DNA strand exchange. One type of lesion requiring BRCA2 for repair is the interstrand-crosslink, which can be induced by chemical reagents such as mitomycin C. The formation of BRCA2 foci after mitomycin C treatment is correlated with the observation that BRCA2 becomes more immobile and suggests that induction of DNA damage causes a redistribution of BRCA2 to DNA damage sites (14). In this study, we use a previously generated mouse embryonic stem (ES) cell line, in which endogenous BRCA2 is homozygously tagged with a HaloTag by knock-in mediated by CRISPR / Cas9. This enables direct, real-time visualization of individual BRCA2 molecules in living cells. Here, we used this cell line (as described in Paul et al., 2021) to investigate the mobility of BRCA2 at damage sites by applying spot-FRAP at BRCA2-HaloTag mitomycin C-induced DNA repair foci. These experiments show a large immobile fraction of BRCA2 at the damage sites, with a 44% fluorescence recovery of the prebleached signal during 50 seconds of data acquisition, corrected for the residual fluorescence signal after photobleaching. (**Figure 1A**). This large immobile fraction for BRCA2 is not observed in the nucleoplasm outside the foci, although the low concentration of BRCA2 in the cell nucleus makes these measurements challenging. Together, our FRAP experiment suggests that upon induction of DNA damage, part of the BRCA2 molecules remain associated to DNA damage sites.

### 3D single-molecule tracking reveals BRCA2 immobilization at individual DNA repair foci

Although FRAP is a powerful technique for studying the dynamic exchange of proteins in cells, it has limited possibilities to differentiate between different types of diffusive behavior of proteins at the single-molecule level, which can be better investigated with single-molecule tracking (15). In this study, we applied 3D single-molecule tracking in the entire nucleus using a multicolor aberration-corrected multifocus microscope (acMFM) (16) (**Figure 1B, Movie S1**). Single-plane single-molecule tracking is restricted by the imaging volume, since molecules can move in and out of the focal plane. Although the axial excitation thickness for HiLo illumination acquisition is in the range of 1-2 µm, the cell nucleus is about 5 µm thick. Furthermore, DNA repair foci, in which BRCA2 dynamics needs to be quantitated, are distributed across the entire nuclear volume, and therefore it is not possible to capture all foci in one focal plane, and hence some of the foci will be out of focus. Finally, both cell nuclei and DNA repair foci are dynamic in their composition and spatial location within the nucleus (17, 18); therefore the regions from which data should be recorded are moving during image acquisition.

Using an acMFM microscope, nine focal planes are projected (approximately 400 nm spacing) on a single camera chip. This allows for rapid instantaneous 3D imaging at 9 planes. To identify DNA damage, EGFP-tr53BP1 was introduced into cells as a marker of DNA breaks. This construct expressing a fragment of the 53BP1 protein (a.a. 1221-1709) is a useful marker for DNA breaks and was previously validated to not interfere with the repair process (19–21). Immunofluorescence experiments confirmed the proper localization of the tr53BP1 protein in our cells, overlapping with the signal of endogenous 53BP1. Upon treatment with the DNA crosslinking agent mitomycin C, tr53BP1 foci can be observed that co-localize with BRCA2 accumulations (**Figure S1**). By applying 3D single-molecule tracking and simultaneously following the foci distribution in cells, we directly correlate the diffusive behavior of BRCA2 with regions of DNA damage (**Figure 1C**). This developed image analysis routine allows us to identify whenever a single BRCA2 molecule is within a region of damage during image acquisition. Subsequently, we tracked the localized molecules in 3D and compared their 2D and 3D displacements inside and outside the mask (**Figure 1D**). This comparison of displacement distributions shows a distribution with shorter displacements inside the mask compared to outside of the mask (**Figure 1E,F**). To validate the observed differences in the displacements between inside and outside the DNA repair foci, we also analyzed the data after randomly redistributing the segmented foci objects along the x- and y-axes of the image plane (see Methods section for details). Compared to the original data, the randomized data shows no difference in the distribution of displacements inside compared to outside the foci (**Figure 1E,F** bottom panels). Additionally, we quantified the number of localizations in the mask and observed that 9.9% of the localizations are within the mask whereas the randomly defined mask contains 1.3% of the localizations. This shows that there is a 7.9 times enrichment of localizations in the data compared to the random mask, confirming the observation that BRCA2 accumulates at foci containing 53BP1 (**Figure 1G**).

Using the described approach, we investigated the motion patterns of BRCA2 specifically at DNA damage. We define molecules to be inside of the focus if they reside at least one frame (50 ms) inside the 53BP1 defined focus volume. This definition allows for comparison of tracked molecules that are inside or in close proximity of the foci, with tracks outside the foci, without introducing edge effects caused by using a binary mask: immobile molecules will have a higher chance of staying within the mask for a longer time and hence will be identified more frequently if we would do the analysis where the entire track needs to be inside the mask. Since we have previously observed that BRCA2 molecules can frequently switch between different diffusive states, we applied a recently developed deep learning approach method to our single-molecule tracking data to segment tracks into tracklets (22). With this method we could define three different diffusive states for BRCA2: fast (*D_app_*=0.44 µm^2^/s), slow (*D_app_*=0.05 µm^2^/s) and immobile (*D_app_*=0.01 µm^2^/s), comparable to previous studies (8, 14, 22). For each individual tracklet (see **Figure S2** for the distribution of track lengths), we fitted the mean square displacement (MSD) to a power-law equation (MSD *∼ t^α^*) permitting the extraction of the anomalous exponent *α*, a metric that gives an indication of the diffusion mode of the track with 0 < *α* < 1 showing specifically subdiffusion, and 1 < *α* < 2 superdiffusion. All molecules exhibited an anomalous exponent *α* below 1 suggesting subdiffusion. Furthermore, both immobile (*α*=0.34) and slow molecules (*α*=0.60) display more subdiffusive motion compared to fast molecules (*α*=0.92) (**Figure 2A**). When comparing different MSD curves for each diffusive state, we observe a clear separation between the states, indicating three distinct mobility patterns. This observation is confirmed by the theoretical MSD curves plotted using the median values in **Figure 2B**.

**Figure 2:**
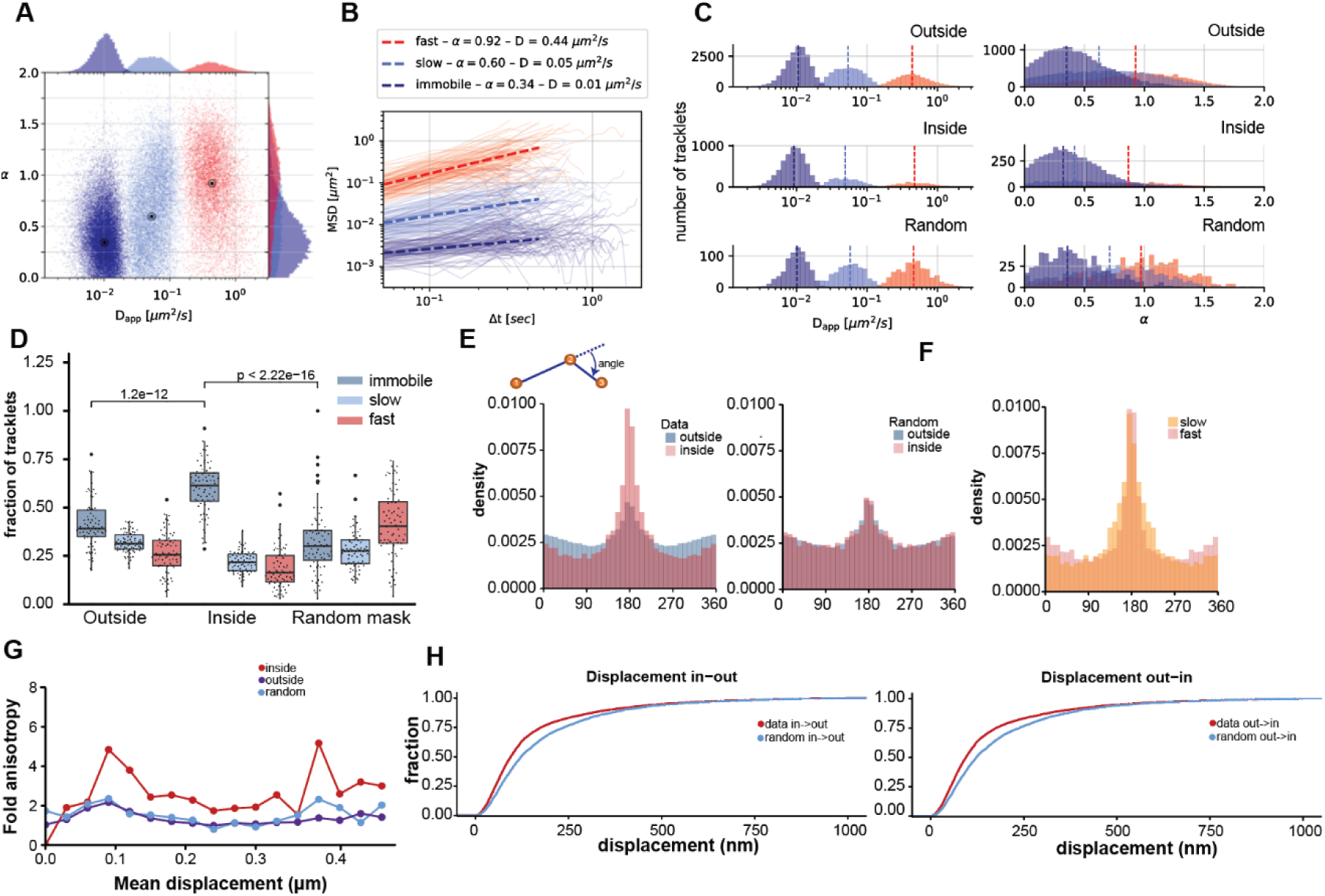
In-depth analysis of mobility in and outside the foci. **A** Scatter plot showing the estimated apparent diffusion constant (Dapp) and anomalous exponent (α) distribution of immobile (blue, Dapp: 0.01 µm2/s, α: 0.34), slow (light blue, Dapp: 0.053, α: 0.60) and fast (red, Dapp: 0.43, α: 0.93) tracklets of BRCA2-HaloTag in mitomycin C treated cells. The black points in the plot indicate the median values for each cluster. **B** For each diffusive state, example of 100 MSD lines with the MSD fit using the power law equation and the medians Dapp and α values. **C** The distribution of Dapp and α values per tracklet outside (Dapp: 0.43;0.05;0.01 µm2/s, α: 0.93;0.62;0.35) , inside (Dapp: 0.42;0.05;0.01 µm2/s, α: 0.87;0.41;0.32) and inside the random mask (Dapp: 0.46;0.06;0.01 µm2/s, α: 0.97;0.71;0.36). The histograms display the overall distribution of all cells imaged. **D** Quantification of the distribution of tracklets outside, inside and inside random mask. The indicated fraction is estimated from the mean fraction per cell. **E** Angular distributions calculated between subsequent localizations. Displacements of immobile molecules are excluded from the analysis. The determined distribution of angles for BRCA2 is plotted for all molecules outside and inside the damage site and compared to a randomized mask. **F** Angular distribution of fast and slow-moving molecules **G** Distribution of the fold anisotropy (ratio between backward angle (180 +/- 15 degrees) versus forward angle (0 +/- 15 degrees) plotted against mean displacement of the track. **H** Cumulative distribution of the displacements of the track segments that transition from inside to outside (left) and from outside to inside (right) of the foci, where immobile molecules are excluded from the distribution. Analysis in this figure is done in 2D using the x-y coordinates in the tracking data. The data consists of four independent experiments of 73 cells in total. p-values are calculated from an unpaired 2-sample Wilcox rank sum test.

Subsequently we compared the diffusion constant *D_app_*and the anomalous exponent *α* of the tracklets in and outside the foci. The diffusion constant of the different fractions is not affected (**Figure 2C**). However, the measured anomalous exponent α is reduced for all fractions within the focus. Quantification of the fraction of tracklets inside the mask shows a clear enrichment in the amount of BRCA2 tracklets in the immobile state at the identified DNA damage sites (42 to 60%), while both the slow (32 to 22%) and fast fraction (26 to 19%) did decrease (**Figure 2D**).

Finally, we compared the results of cells treated with mitomycin C with untreated cells (**Figure S3**). In untreated mouse ES cells, spontaneous foci can be observed and immobilization of BRCA2 has also been observed under unchallenged conditions, although to a lesser extent. We confirmed these observations in our current experiments, where in the untreated cells we see that the immobile fraction inside (52% versus 60%) is lower while the immobile fraction outside the foci is comparable (38% versus 39%) to the treated cells. The median diffusion constant of the different fractions was not different compared to the cells treated with mitomycin C. Similarly, to treated cells the value of α was reduced for diffusing fractions inside the foci (**Figure 2, S3**). Interestingly, also the fraction of BRCA2 molecules that resides in 53BP1 foci is lower compared to treated cells (3.4% versus 9.9%), indicating that at spontaneous foci less BRCA2 molecules accumulate.

### Diffusive BRCA2 molecules exhibit enhanced subdiffusive motion at DNA repair foci

Our aim was to further investigate the mechanism by which BRCA2 accumulates at the damage sites. Tracking analysis showed that at sites of DNA damage, next to an increased immobile fraction, slow diffusing molecules exhibited subdiffusive diffusion (alpha<1). An alternative approach to study the subdiffusion of proteins in cells is to look at the angular distribution, a sign of anisotropic motion of the tracked molecules. Molecules that exhibit regular diffusion will move in a random direction independent of their previous position, but when motion is confined, molecules are more likely to make a backward step than forward with respect to the previous step (5, 7). To investigate only diffusive molecules, we excluded tracklets that are assigned by our state segmentation method as immobile from the analysis. In fact, after calculating the angles, mobile BRCA2 molecules appear to have an anisotropic motion pattern, and this anisotropy is enhanced at the DNA repair foci (**Figure 2E**). This is the case for both slow and fast diffusing molecules, although the fastest diffusing molecules show this behavior to a lesser extent (**Figure 2F**). Subsequently we quantified the fold increase in the backward angle (180 +/- 15 degrees) versus the forward angle (0 +/- 15 degrees) compared to the average displacement of the tracked molecule at that time (**Figure 2G**). We can observe a peak between 80 and 150 nm of the average displacement, which suggests that, especially molecules having shorter displacements, are, transiently, more confined to a subregion of the focus. Then we quantified the distribution of displacements of molecules that transition from inside to outside or vice-versa. We observe that for both transitions the smaller displacements are observed compared to the random mask (**Figure 2H**). However, the transitions in either direction appear similar for both the data set and the random mask, suggesting that movement in and out of the focus is not restricted.

To evaluate whether the BRCA2 molecules inside the foci do display confined motion within the repair focus, we fitted the mean-square-displacement data to a confined motion model which estimates a radius of confinement *R_c_*. For all immobile tracklets localized inside or outside the mask, we estimated the plateau of each averaged MSD line (**Figure S4**). The results show an estimated *R_c_* of about 0.126 µm outside the foci and 0.144 µm inside the foci (versus 0.136 µm inside the randomly generated mask). We reproduced the same measurements for the moving molecules (slow and fast combined), although the motion is less confined regarding the previously calculated *α* values, leading to a potential delayed MSD plateau. We found a *R_c_* value of 1.57 µm outside and surprisingly a very large Rc value of about 6.21 µm inside the foci (versus 2.70 µm inside the random masks). Although the curve fits well to the data, the lack of a clear plateau in the MSD curve suggests that, for the slow and fast diffusing molecules, the molecules do not experience confinement correlating with focus size. For comparison we also fitted the same MSD curves to an anomalous diffusion model which displays a non-linear relation between displacement and time, but without constraint or a boundary (**Figure S5**). This also resulted in a good fit of the data suggesting that the motion of BRCA2 at repair foci can be explained by transient binding of molecules within the damage site resulting in the observed anomalous diffusion patterns.

### BRCA2 is localized in multiple clusters of several BRCA2-molecules at DNA repair foci

To better understand the organization of BRCA2 in the foci, we have applied dSTORM imaging of BRCA2-Halo to visualize the spatial arrangement of BRCA2 under similar experimental conditions in 53BP1 foci after mitomycin C treatment. Previous experiments in human cells using anti-BRCA2 antibodies showed the presence of multiple BRCA2 clusters per focus (23). By using direct HaloTag labelling of BRCA2-Halo with Alexa647-HaloTag ligand, we can now more accurately localize BRCA2 in foci by single-molecule localization microscopy. We used 53BP1 foci to identify regions of interest to determine the localization of BRCA2 within. Visual comparison of the localization of 53BP1 and BRCA2 in the foci indicates that BRCA2 is frequently excluded from regions that are dense in 53BP1 signal (**Figure 3A**). Within the 53BP1 foci we quantified BRCA2 to localize in multiple small clusters of about 50 nm in size (**Figure 3A-C**). Furthermore, we quantified the distance between BRCA2 clusters (Figure 3D). This analysis shows that most BRCA2 clusters are most frequently 200 nm spaced apart with respect to their centre of mass. To confirm that BRCA2 cluster formation was present in cells in lateS/G2, where BRCA2 is expected to contribute to homology-directed repair, we applied dSTORM imaging in cells that were labeled shortly with EdU prior to fixation (**Figure S6**). In addition to the differential location between BRCA2 and 53BP1, we also identified that RPA is differentially localized in the repair foci with respect to BRCA2 (**Figure S7**).

**Figure 3:**
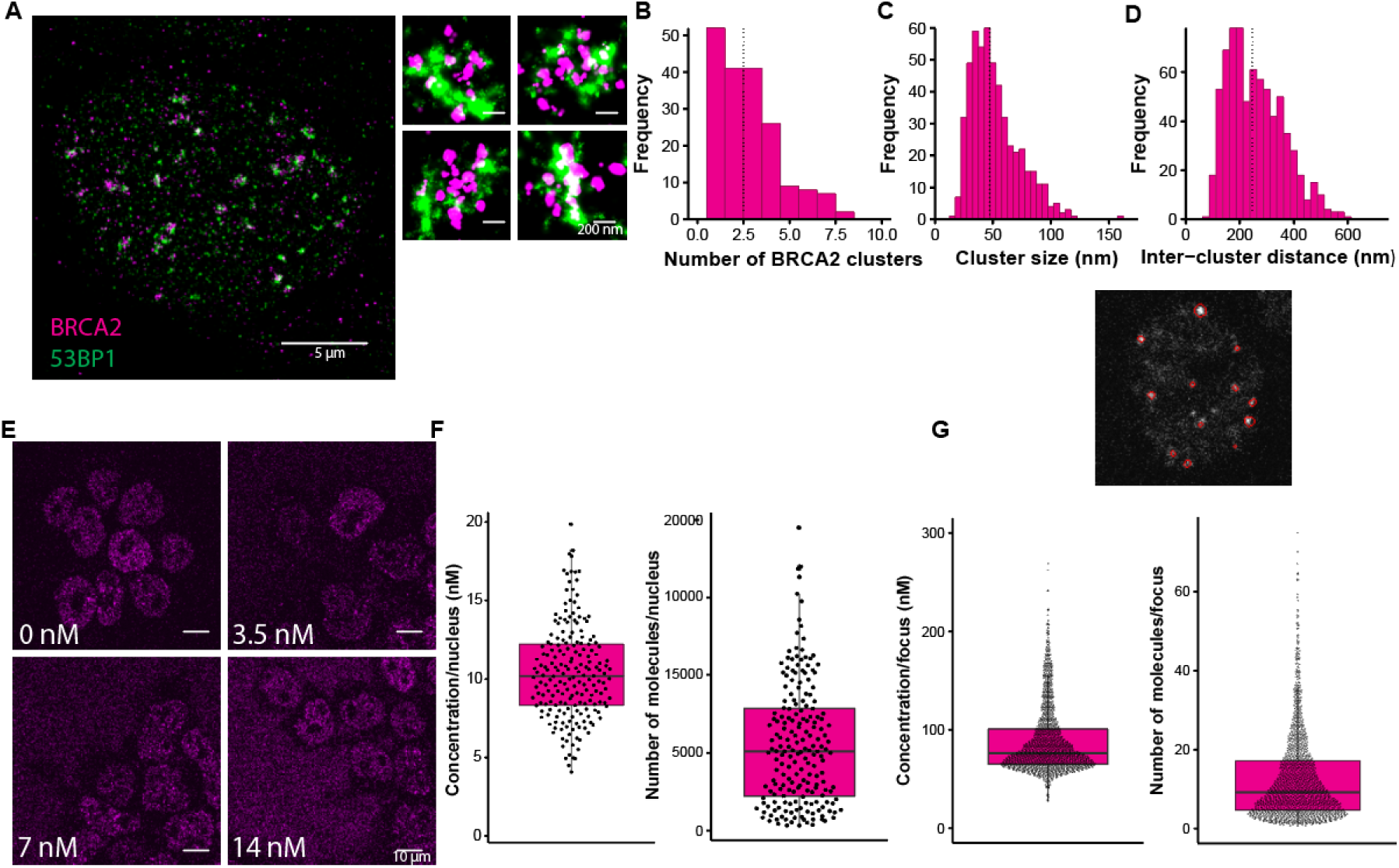
Nanoscale localization of BRCA2 in MMC-induced DNA repair foci and quantification of nuclear BRCA2 concentration. **A** Two-color dSTORM image of BRCA2-Halo::Alexa647 and 53BP1 (indirect immunofluorescence CF568). Typical examples of foci with both BRCA2 and 53bp1 signal. **B** Quantification of the number of BRCA2 clusters within individual automatically segmented 53BP1 foci. **C-D** Quantification of the size of the respective clusters and the centre-to-centre distance between the clusters within the individual foci. Data from 7 cells in total from two imaging sessions. Analysis of 7 images with a total of 189 foci analyzed. Dashed line indicate the median of the distributions. **E** The cellular concentration of BRCA2 was estimated by titration of increasing concentrations of the labeled GST-HaloTag protein in the cell medium. The voxel intensity was used to estimate the local concentration. **F** The nuclear concentration was estimated by 3D segmentation of the cell nuclei. The nuclear concentration is defined as the average nuclear concentration within all voxels in the 3D nuclear volume, whereas the number of BRCA2 molecules per nucleus is calculated by multiplying the concentration with the estimated nuclear volume. **G** Within the identified nuclei, the foci were segmented in 3D. The image shows an example of the segmentation in a single confocal slice. The plots show the results of one replicate, results from additional replicates (n=3) are summarized in Fig. S6.

We validated that the inhomogeneous localization that we observed by dSTORM is unlikely to be an artifact caused by repetitive photoblinking, which is known to cause apparent clustering in dSTORM imaging. First we used an algorithm to correct for blinking artifacts and reconstruct the true emitters in a few of the foci (24). This showed that after blinking correction the BRCA2 clusters are still visible (**Figure S8**). Furthermore, we also used 2D-STED imaging of BRCA2 and 53BP1 to determine the super-resolution localization of BRCA2. While the lateral resolution of STED is lower, hence less details are visible it is not affected by blinking artifacts like in dSTORM. With STED imaging we also observe that molecules are differentially organized and that BRCA2 can form multiple clusters per 53BP1 focus (**Figure S8**).

Subsequently, we wanted to estimate the concentration of BRCA2 in mouse ES cell nuclei to correlate the amount of BRCA2 present at repair foci with the clusters that are present in the foci. For this purpose, we imaged live BRCA2-HaloTag cells by confocal microscopy, while titrating known concentrations of JF646-labeled GST-HaloTag protein in cell culture medium. This allowed us to determine the correlation between voxel intensity and HaloTag protein concentration in the images (**Figure 3E, Figure S9**). Although we did not estimate the actual excitation confocal volume of a voxel, we were able to determine the local concentration of BRCA2-Halo per pixel. With the assumption that the nuclei and foci are larger than the confocal volume, we can use the integrated intensity of the objects to estimate the local concentration and total number of BRCA2 molecules in individual nuclei. This shows a nuclear concentration of 10 nM or about 5,000 molecules per cell nucleus (**Figure 3F**). Additionally, we were able to segment individual BRCA2 foci to estimate the number of BRCA2 molecules per focus. This analysis indicates that the local concentration of BRCA2 is nearly 10 times higher within repair foci (∼100 nM) whereas the mean number of BRCA2 molecules per focus was estimated at approximately 25 (**Figure 3G**). This increase in concentration of BRCA2 molecules at the repair foci is comparable to increased number of localizations within repair foci we observed with single-molecule tracking (**Figure 1G**). The concentration of BRCA2 we determined in cells is consistent with the quantification of the amount of cellular BRCA2-Halo protein labelled with JF646 and on SDS-PAGE, while using the GST-HaloTag protein with defined concentration as reference (**Figure S9**). Quantification of the total intensity of the protein bands normalized to the number of cells in the sample loaded, reports the average number of BRCA2 molecules per nucleus.

## DISCUSSION

In this study we have investigated the dynamic behavior of BRCA2 specifically at 53BP1 foci induced by the DNA crosslinker mitomycin C. Instantaneous multiplane two-color imaging allowed us to detect the BRCA2 molecules localized in DNA repair foci and study their mobility. Previous single-molecule tracking studies of BRCA2 have shown an increase in the immobile fraction of BRCA2 after DNA damage induction (8, 14). Here, we show that this immobilization is caused by BRCA2 that is localized to DNA repair foci. We identified that only about 10% of the BRCA2 molecules that were tracked within the nucleus were accumulated at the foci, which explains the subtle increase in the immobilization of BRCA2 measured throughout the nucleus, as we have previously observed. In depth-analysis of the 3D tracking data revealed that next to the increased immobilization, diffusing BRCA2 molecules inside the foci show increased anomalous diffusion. This anomalous diffusion is most evident in slow moving BRCA2 molecules, with an increased confinement for molecules moving between 80 and 150 nm in a 50 ms time interval. A similar pattern of anomalous diffusion is observed for BRCA2 molecules outside the foci, but to a lesser extent (Figure 2E-G). Similar motion patterns have been observed for other nuclear proteins such as CTCF and transcription factors (Sox2 and Oct4) (6, 7, 27), which contribute to the function of these proteins, such as improved target search in chromatin. For BRCA2 complexes, this could be a mechanism to efficiently probe the chromatin environment for the presence of DNA damage. The difference in the chromatin environment (*e.g.,* changes in specific histone modifications) present at DNA damage sites could enhance transient binding of BRCA2 complexes at the damaged chromatin. This will allow BRCA2, even at its low concentration, to dynamically associate with the damaged genomic region. This local difference in affinity could serve as a mechanism to retain and concentrate BRCA2 molecules at DNA damage for a longer period and could explain the motion patterns observed for BRCA2.

A similar mechanism could operate within a focus. Our observation by single-molecule localization microscopy (**Figure 3**) of BRCA2 clusters within a focus suggests that the chromatin environment within a focus is not homogeneous. The measured distance between these intra-focus BRCA2 clusters (**Figure 3**) is consistent with the possibility of displacement of BRCA2 molecules between these clusters (resulting in the class of BCRA2 molecules with confined motion), while molecules with larger displacements would lose their association with the clusters. It remains to be investigated what interactions drive these motion patterns, but it should be considered that BRCA2 is present in complex with several other repair proteins such as PALB2 and RAD51. For example, loss of the PALB2-BRCA2 interaction prevents BRCA2 localization in foci (9). On the contrary, the accumulation of BRCA2 does not depend on its highly conserved DNA binding domain and the C-terminal domain (8), which, at least in vitro, interact with (single-stranded) DNA (28). An explanation is that BRCA2 is bound to chromatin surrounding the DNA damage mainly through its N-terminal interaction with PALB2, without the requirement of all functional domains required to promote all BRCA2 functions. In this way, the accumulation of BRCA2 is uncoupled from its canonical function of loading RAD51 onto RPA coated DNA. On the other hand, the transient interactions will help BRCA2 to reside at a focus for longer, that might be required to give BRCA2 sufficient time to exert its function of regulating RAD51 filament formation.

### Mechanisms of accumulation of BRCA2 at DNA lesions

The dynamics of a larger number of different DNA repair proteins has been investigated in living cells. It shows that while many of these proteins accumulate at sites of DNA damage, they can do so through different mechanisms (29). While RAD51 and RPA are stably bound (17, 30), RAD54, another protein in the same DNA repair pathway, shows a different dynamic behaviour (17). Interestingly, while RAD54 visually accumulates in foci, immobilization of the protein is hardly observed, unless its ATP activity is affected (31). In recent years, liquid-liquid phase separation (LLPS) has been proposed as an additional mechanism to drive compartmentalization of proteins in DNA repair foci, as well as in other nuclear and cellular processes. 53BP1 is one of the DNA repair factors that forms accumulations with liquid-like characteristics (25, 26). Furthermore, RAD52, a RAD51 mediator in yeast, can form condensates *in vitro* and *in vivo* (32). Another study using single-molecule tracking showed that RAD52 in budding yeast shows reduced and confined mobility in foci, while the RPA subunit Rfa1, shows clear immobilization (30). To drive the formation of condensates proteins need to be present at a high enough concentration, however BRCA2 is present at a low concentration in the cell nucleus and within the foci only tens of BRCA2 molecules are present (**Figure 3**). This suggests that BRCA2 by itself is not phase separated but is trapped in DNA lesion-containing compartments where it undergoes transient immobilization through interaction with other proteins or DNA in the foci. As the examples above show, repair factors can accumulate through different mechanisms of which transient binding and the formation of phase separated compartments are the most well-described mechanisms. Differentiating between these different modes of accumulation is not straightforward and many different parameters have been proposed to prove the formation of condensates through liquid-liquid phase separation (33). With the current data on BRCA2 mobility we are not able to apply modelling of the different models as we lack accurate information on the strength and reversibility of the binding of BRCA2 within the foci, nor do we have an accurate estimation of the surface potential in a phase-separation model (34).

Through different approaches of analysing our tracking data we do observe that the diffusion rate of the diffusive fraction of BRCA2 is not affected inside the repair focus. If BRCA2 would be accumulating through interactions driven by the presence of phase separation, a change in the measured diffusion rate of the diffusing molecules due to transient molecular interactions would be expected, as has been observed for RAD52 in yeast. Simultaneously we observe an increase in subdiffusive motion in the repair foci, however using the mean-square displacement curve of the diffusive molecules we are not able to observe a clear plateau in the MSD curve and the fitted confinement radius does not match the size of repair foci. This suggests that the mobility of BRCA2 is not restricted by the boundary of the focus, but that transient binding contributes to the accumulation of BRCA2.

### Dynamic organization of BRCA2 at DNA repair foci

In line with our previous publication on the localization of BRCA2 using antibodies, BRCA2-HaloTag is localized in small clusters at DNA break sites in fixed cells. Quantification of the amount of BRCA2 in cell nuclei and repair foci specifically shows that only tens of BRCA2 molecules are present in DNA repair foci; therefore, individual clusters of BRCA2 within a focus will contain only a few BRCA2 molecules (**Figure 3**). The emerging picture from several studies that use super resolution microscopy to study the architecture of DNA repair foci is that proteins organized in repair foci in specific patterns rather than a homogeneous mix of different repair factors (35–37). For example, by single-molecule localization microscopy, we have observed that part of the RAD51 proteins that are present at repair foci do not localize with BRCA2 (23). It is interesting to consider that even for proteins that are suggested to accumulate by LLPS, such as 53BP1, there is still spatial inhomogeneity of the protein within the focus, instead of a homogeneous distribution of protein. A challenge for future studies will be to converge the observed mobility patterns of the different proteins with their nanoscale localization at foci. An important aspect that still needs to be addressed is how dynamic the overall organization of repair foci is over time. Future studies using super-resolution microscopy in living cells using methods to enhance spatial and temporal resolution, will hopefully allow to further delineate the significance of spatiotemporal dynamics of DNA repair foci that protect the cells against genomic instability.

## DATA AVAILABILITY

Source data is deposited at the Zenodo repository. Original images of the tracking experiments can be found at: http://doi.org/10.5281/zenodo.10144073 Source data of other experiments can found at: http://doi.org/10.5281/zenodo.8143173

## AUTHOR CONTRIBUTIONS

MWP, JA, TLC, RK, and CW designed research; MWP, AT and JA performed research; MWP, JA, EW, HK, RvG and IS contributed new reagents/analytic tools; MWP, JA, EW, HK, RvG, and IS analysed data; MWP, RK and CW wrote the paper. All authors discussed the results and commented on the manuscript.

## ACKNOWLEDGEMENTS

The authors would like to dedicate this work to our friend and colleague Claire Wyman, who passed away during completion of this research. Aberration corrected multifocal microscopy (acMFM) was performed in collaboration with the Advanced Imaging Center at Janelia Research Campus, a facility jointly supported by the Gordon and Betty Moore Foundation and the Howard Hughes Medical Institute. We thank the Optical Imaging Centre at the Erasmus MC for use and technical assistance with the optical microscopes. We thank the Josephine Nefkens Cancer Program for infrastructure support.

## REFERENCES

1. Paul, M.W., Sidhu, A., Liang, Y., van Rossum-Fikkert, S.E., Odijk, H., Zelensky, A.N., Kanaar, R. and Wyman, C. (2021) Role of BRCA2 DNA-binding and C-terminal domain on its mobility and conformation in DNA repair. eLife, 10, e67926.

2. Reuter, M., Zelensky, A., Smal, I., Meijering, E., van Cappellen, W.A., de Gruiter, H.M., van Belle, G.J., van Royen, M.E., Houtsmuller, A.B., Essers, J., et al. (2014) BRCA2 diffuses as oligomeric clusters with RAD51 and changes mobility after DNA damage in live cells. The Journal of Cell Biology, 207, 599–613.

3. Zelensky, A.N., Schimmel, J., Kool, H., Kanaar, R. and Tijsterman, M. (2017) Inactivation of Pol θ and C-NHEJ eliminates off-target integration of exogenous DNA. Nature Communications, 8.

4. Yang, K.S., Kohler, R.H., Landon, M., Giedt, R. and Weissleder, R. (2015) Single cell resolution in vivo imaging of DNA damage following PARP inhibition. Scientific Reports, 5, 10129.

5. Abrahamsson, S., Chen, J., Hajj, B., Stallinga, S., Katsov, A.Y., Wisniewski, J., Mizuguchi, G., Soule, P., Mueller, F., Darzacq, C.D., et al. (2013) Fast multicolor 3D imaging using aberration-corrected multifocus microscopy. Nature Methods, 10, 60–63.

6. Bohrer, C.H., Yang, X., Thakur, S., Weng, X., Tenner, B., McQuillen, R., Ross, B., Wooten, M., Chen, X., Zhang, J., et al. (2021) A pairwise distance distribution correction (DDC) algorithm to eliminate blinking-caused artifacts in SMLM. Nat Methods, 18, 669–677.

7. Sbalzarini, I.F. and Koumoutsakos, P. (2005) Feature point tracking and trajectory analysis for video imaging in cell biology. J. Struct. Biol., 151, 182–195.

8. Arts, M., Smal, I., Paul, M.W., Wyman, C. and Meijering, E. (2019) Particle Mobility Analysis Using Deep Learning and the Moment Scaling Spectrum. Sci Rep, 9, 1–10.

9. Ushey, K., Allaire, J.J. and Tang, Y. (2019) reticulate: Interface to ‘Python’.

10. Metzler, R., Jeon, J.-H., Cherstvy, A.G. and Barkai, E. (2014) Anomalous diffusion models and their properties: non-stationarity, non-ergodicity, and ageing at the centenary of single particle tracking. Phys. Chem. Chem. Phys., 16, 24128–24164.

11. Lerner, J., Gómez-García, P.A., McCarthy, R.L., Liu, Z., Lakadamyali, M. and Zaret, K.S. (2020) Two-parameter single-molecule analysis for measurement of chromatin mobility. STAR Protocols, 1, 100223.

12. Ershov, D., Phan, M.-S., Pylvänäinen, J.W., Rigaud, S.U., Le Blanc, L., Charles-Orszag, A., Conway, J.R.W., Laine, R.F., Roy, N.H., Bonazzi, D., et al. (2022) TrackMate 7: integrating state-of-the-art segmentation algorithms into tracking pipelines. Nat Methods, 19, 829–832.

13. Pike, J.A., Khan, A.O., Pallini, C., Thomas, S.G., Mund, M., Ries, J., Poulter, N.S. and Styles, I.B. (2020) Topological data analysis quantifies biological nano-structure from single molecule localization microscopy. Bioinformatics, 36, 1614–1621.

14. Paul, M.W., de Gruiter, H.M., Lin, Z., Baarends, W.M., van Cappellen, W.A., Houtsmuller, A.B. and Slotman, J.A. (2019) SMoLR: visualization and analysis of single-molecule localization microscopy data in R. BMC Bioinformatics, 20.

15. Schmidt, U., Weigert, M., Broaddus, C. and Myers, G. (2018) Cell Detection with Star-convex Polygons. arXiv:1806.03535 [cs], 11071, 265–273.

16. McQuin, C., Goodman, A., Chernyshev, V., Kamentsky, L., Cimini, B.A., Karhohs, K.W., Doan, M., Ding, L., Rafelski, S.M., Thirstrup, D., et al. (2018) CellProfiler 3.0: Next-generation image processing for biology. PLOS Biology, 16, e2005970.

17. Aaron, J., Wait, E., DeSantis, M. and Chew, T. (2019) Practical Considerations in Particle and Object Tracking and Analysis. Current Protocols in Cell Biology, 10.1002/cpcb.88.

18. Essers, J., Houtsmuller, A.B., van Veelen, L., Paulusma, C., Nigg, A.L., Pastink, A., Vermeulen, W., Hoeijmakers, J.H.J. and Kanaar, R. (2002) Nuclear dynamics of RAD52 group homologous recombination proteins in response to DNA damage. EMBO J, 21, 2030– 2037.

19. Krawczyk, P.M., Borovski, T., Stap, J., Cijsouw, T., Cate, R. t., Medema, J.P., Kanaar, R., Franken, N.A.P. and Aten, J.A. (2012) Chromatin mobility is increased at sites of DNA double-strand breaks. Journal of Cell Science, 125, 2127–2133.

20. Dimitrova, N., Chen, Y.-C.M., Spector, D.L. and Lange, T. de (2008) 53BP1 promotes non-homologous end joining of telomeres by increasing chromatin mobility. Nature, 456, 524–528.

21. Lottersberger, F., Karssemeijer, R.A., Dimitrova, N. and de Lange, T. (2015) 53BP1 and the LINC Complex Promote Microtubule-Dependent DSB Mobility and DNA Repair. Cell, 163, 880–893.

22. Hansen, A.S., Amitai, A., Cattoglio, C., Tjian, R. and Darzacq, X. (2019) Guided nuclear exploration increases CTCF target search efficiency. Nat Chem Biol, 10.1038/s41589-019-0422-3.

23. Izeddin, I., Récamier, V., Bosanac, L., Cissé, I.I., Boudarene, L., Dugast-Darzacq, C., Proux, F., Bénichou, O., Voituriez, R., Bensaude, O., et al. (2014) Single-molecule tracking in live cells reveals distinct target-search strategies of transcription factors in the nucleus. eLife, 3.

24. Sánchez, H., Paul, M.W., Grosbart, M., van Rossum-Fikkert, S.E., Lebbink, J.H.G., Kanaar, R., Houtsmuller, A.B. and Wyman, C. (2017) Architectural plasticity of human BRCA2– RAD51 complexes in DNA break repair. Nucleic Acids Research, 45, 4507–4518.

25. Kilic, S., Lezaja, A., Gatti, M., Bianco, E., Michelena, J., Imhof, R. and Altmeyer, M. (2019) Phase separation of 53BP1 determines liquid-like behavior of DNA repair compartments. The EMBO Journal, 0, e101379.

26. Pessina, F., Giavazzi, F., Yin, Y., Gioia, U., Vitelli, V., Galbiati, A., Barozzi, S., Garre, M., Oldani, A., Flaus, A., et al. (2019) Functional transcription promoters at DNA double-strand breaks mediate RNA-driven phase separation of damage-response factors. Nat Cell Biol, 21, 1286–1299.

27. Alexander, J.M., Guan, J., Li, B., Maliskova, L., Song, M., Shen, Y., Huang, B., Lomvardas, S. and Weiner, O.D. (2019) Live-cell imaging reveals enhancer-dependent Sox2 transcription in the absence of enhancer proximity. eLife, 8, e41769.

28. Liu, Z., Legant, W.R., Chen, B.-C., Li, L., Grimm, J.B., Lavis, L.D., Betzig, E. and Tjian, R. (2014) 3D imaging of Sox2 enhancer clusters in embryonic stem cells. eLife, 3.

29. Xia, B., Sheng, Q., Nakanishi, K., Ohashi, A., Wu, J., Christ, N., Liu, X., Jasin, M., Couch, F.J. and Livingston, D.M. (2006) Control of BRCA2 Cellular and Clinical Functions by a Nuclear Partner, PALB2. Molecular Cell, 22, 719–729.

30. Yang, H. (2002) BRCA2 Function in DNA Binding and Recombination from a BRCA2-DSS1-ssDNA Structure. Science, 297, 1837–1848.

31. Miné-Hattab, J., Liu, S. and Taddei, A. (2022) Repair Foci as Liquid Phase Separation: Evidence and Limitations. Genes, 13, 1846.

32. Miné-Hattab, J., Heltberg, M., Villemeur, M., Guedj, C., Mora, T., Walczak, A.M., Dahan, M. and Taddei, A. (2021) Single molecule microscopy reveals key physical features of repair foci in living cells. eLife, 10, e60577.

33. Agarwal, S., van Cappellen, W.A., Guénolé, A., Eppink, B., Linsen, S.E.V., Meijering, E., Houtsmuller, A., Kanaar, R. and Essers, J. (2011) ATP-dependent and independent functions of Rad54 in genome maintenance. The Journal of Cell Biology, 192, 735– 750.

34. Oshidari, R., Huang, R., Medghalchi, M., Tse, E.Y.W., Ashgriz, N., Lee, H.O., Wyatt, H. and Mekhail, K. (2020) DNA repair by Rad52 liquid droplets. Nat Commun, 11, 1–8.

35. McSwiggen, D.T., Mir, M., Darzacq, X. and Tjian, R. (2019) Evaluating phase separation in live cells: diagnosis, caveats, and functional consequences. Genes Dev., 10.1101/gad.331520.119.

36. Heltberg, M.L., Miné-Hattab, J., Taddei, A., Walczak, A.M. and Mora, T. (2021) Physical observables to determine the nature of membrane-less cellular sub-compartments. bioRxiv, 10.1101/2021.04.01.438041.

37. Chapman, J.R., Sossick, A.J., Boulton, S.J. and Jackson, S.P. (2012) BRCA1-associated exclusion of 53BP1 from DNA damage sites underlies temporal control of DNA repair. Journal of Cell Science, 125, 3529–3534.

38. Whelan, D.R., Lee, W.T.C., Yin, Y., Ofri, D.M., Bermudez-Hernandez, K., Keegan, S., Fenyo, D. and Rothenberg, E. (2018) Spatiotemporal dynamics of homologous recombination repair at single collapsed replication forks. Nature Communications, 9, 3882.

39. Ochs, F., Karemore, G., Miron, E., Brown, J., Sedlackova, H., Rask, M.-B., Lampe, M., Buckle, V., Schermelleh, L., Lukas, J., et al. (2019) Stabilization of chromatin topology safeguards genome integrity. Nature, 574, 571–574.

