## Supplementary figures for "Distinct mobility patterns of BRCA2 molecules at DNA damage sites"

#### **Supporting Information for**
### Distinct mobility patterns of BRCA2 molecules at DNA damage sites

Maarten W. Paul[1,*], Jesse Aaron[2], Eric Wait[2,3], Romano M. van Genderen[1], Arti Tyagi[1,4], Hélène Kabbech[5], Ihor Smal[1,6], Teng-Leong Chew[2], Roland Kanaar[1] and Claire Wyman[1,7]

Corresponding author: Maarten W. Paul


**This PDF file includes:**

Figures S1 to S9
Legends for Movies S1 to S3

**Other supporting materials for this manuscript include the following:**

Movies S1 to S3

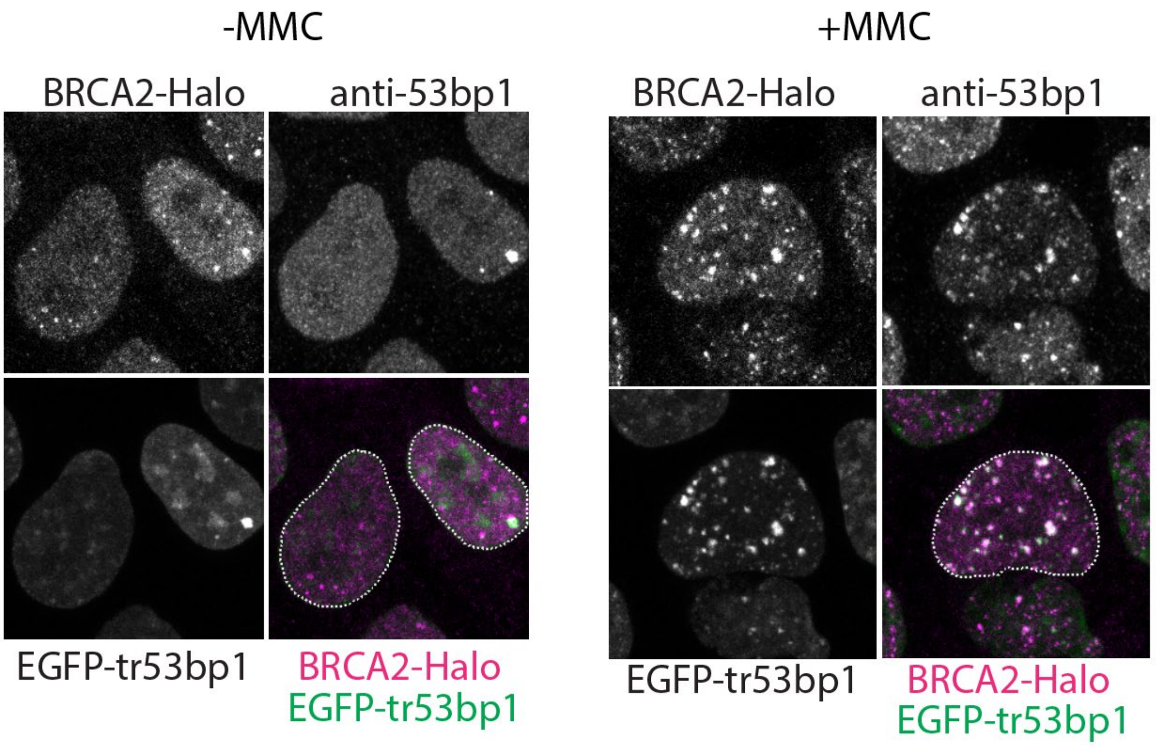


**Fig. S1. Validation of EGFP-tr53bp1 construct.** Immunostaining against endogenous 53bp1 in fixed cells expressing BRCA2-HaloTag and EGFP-tr53bp1 without and with treatment with mitomycin C (MMC)

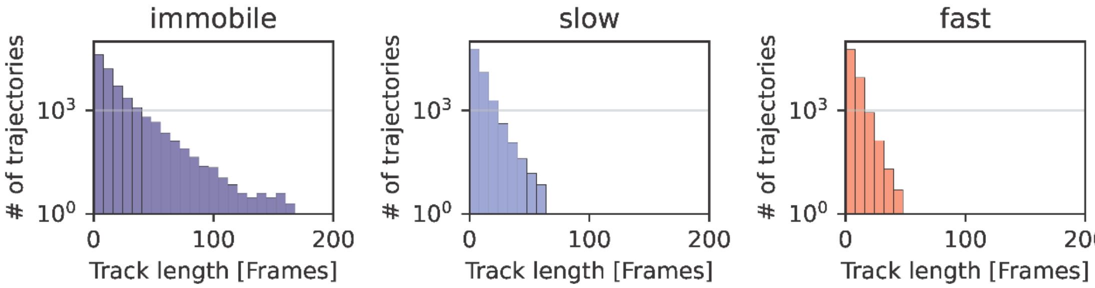


**Fig. S2.** Distribution of track lengths for the different fractions of BRCA2-HaloTag of the dataset of cells treated with mitomycin C. A total of 516 813 immobile, 251 161 slow and 204 666 fast tracklets are in the data set.

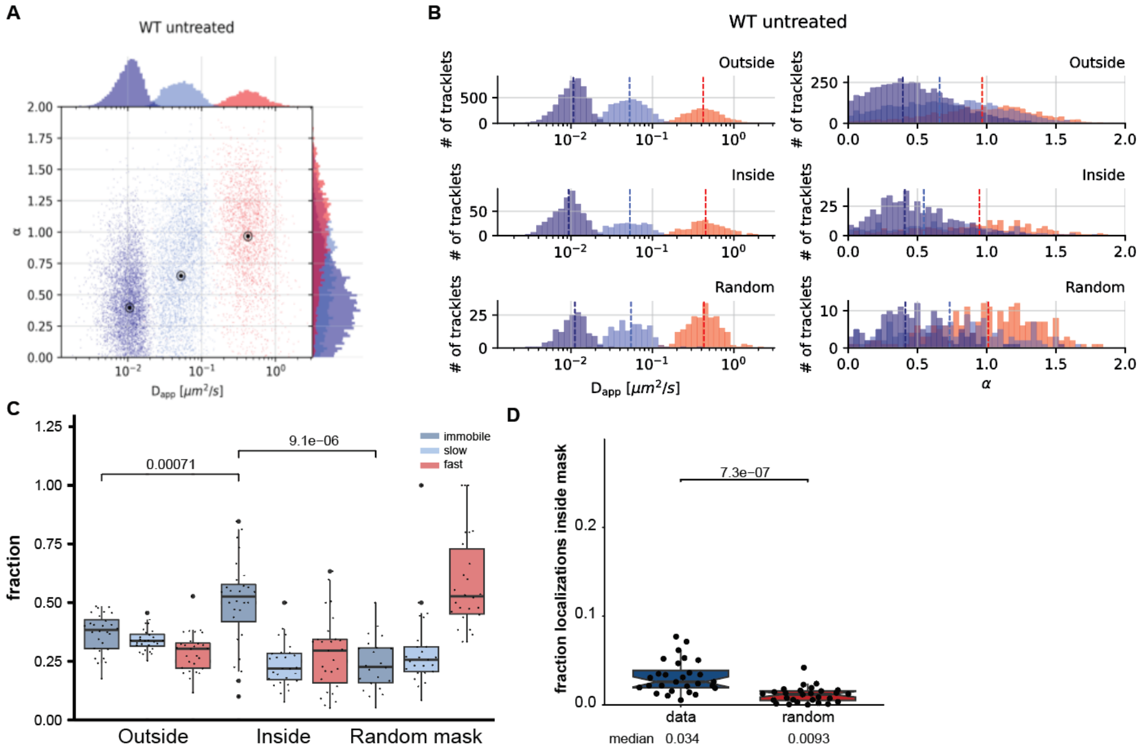


s

**Fig S3.** Analysis results for untreated BRCA2-HaloTag. Analysis was performed the same way as for the mitomycin C treated cells in Figure and Figure 2. **A** Scatter plot showing the estimated apparent diffusion constant ($D_{app}$) and anomalous exponent ($\alpha$) distribution of immobile (blue, $D_{app}$: 0.01 μm$^2$/s, $\alpha$: 0.40), slow (light blue, $D_{app}$: 0.052, $\alpha$: 0.65) and fast (red, $D_{app}$: 0.42, $\alpha$: 0.97) tracklets of BRCA2-HaloTag in untreated cells. Black points in the plot indicate median values for each cluster. **B** The distribution of $D_{app}$ and $\alpha$ values per tracklet outside ($D_{app}$: 0.43;0.05;0.01 μm$^2$/s, $\alpha$: 0.97;0.62;0.35) , inside ($D_{app}$: 0.45;0.05;0.01 μm$^2$/s, $\alpha$: 0.95;0.55;0.41) and inside the random mask ($D_{app}$: 0.43;0.06;0.01 μm$^2$/s, $\alpha$: 1.01;0.73;0.41). The histograms display the overall distribution of all cells imaged. **C** Quantification of the distribution of tracklets outside, inside and inside random mask. The distribution of fractions is measured from the mean fractions per cell. **D** Fraction of localizations in and outside the defined mask for the data compared to the random mask. The total number of 28 cells were analyzed in two independent experiments.

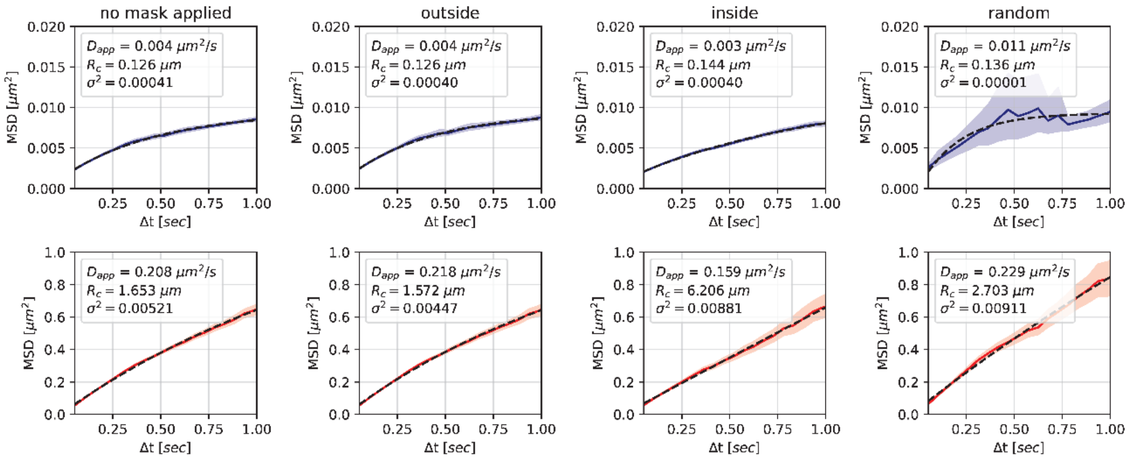


**Fig S4.** Fitting of MSD curves with radius of confinement $R_c$ for the immobile fraction (above) and combined slow and mobile (below) fractions for all tracks (no mask), outside, inside tracks and with a random mask. The MSD curve of the immobile fraction in the random mask looks more noisy due to the lower number of tracks in the immobile state in the random mask.

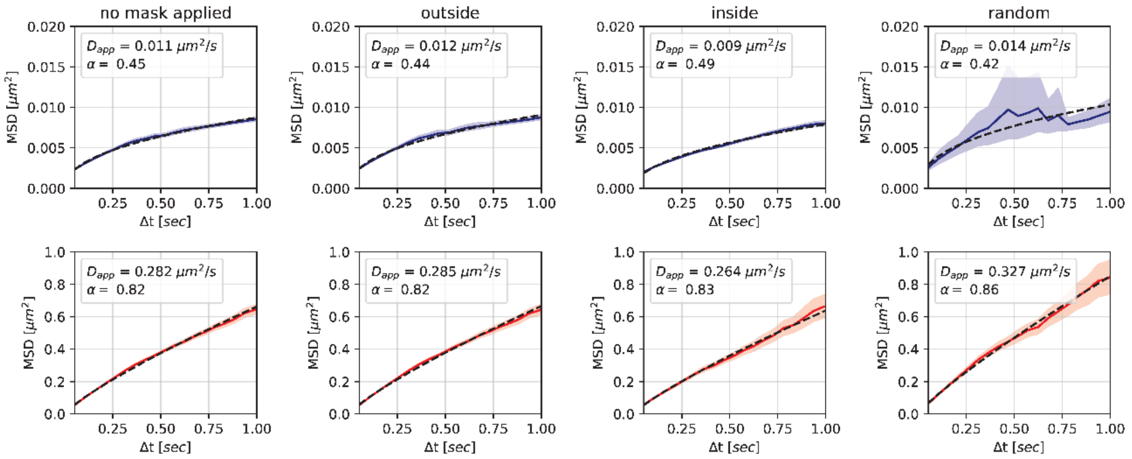


**Fig S5.** Fitting of MSD curves with $D_{app}$ and anomalous exponent $\alpha$ for the immobile fraction (above) and combined slow and mobile (below) fractions for all tracks (no mask), outside, inside tracks and with a random mask. The MSD curve of the immobile fraction in the random mask looks more noisy due to the lower number of tracks in the immobile state in the random mask.

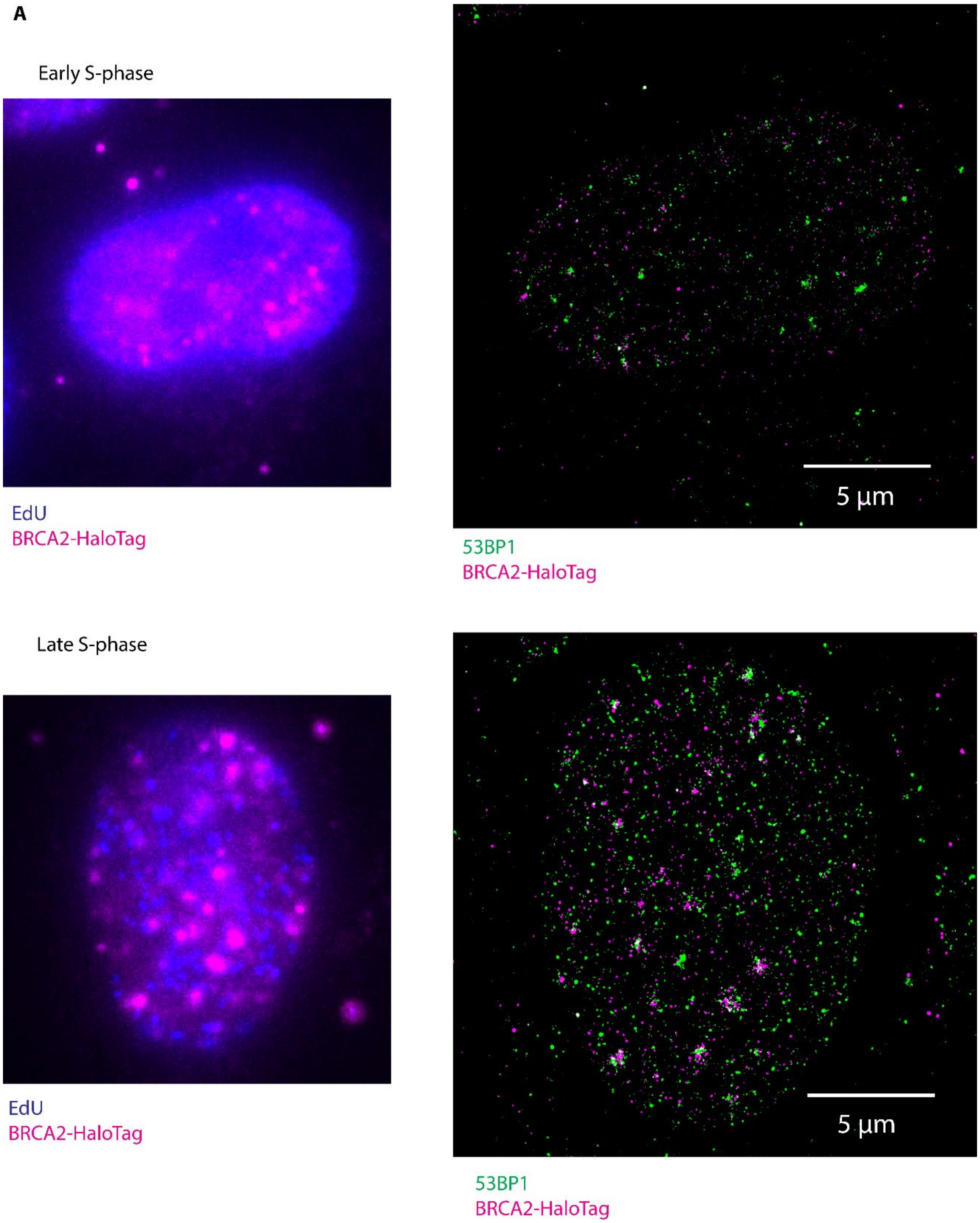


**Fig S6**. dSTORM image of an early and late-S phase cell identified by EdU staining (blue) after mitomycin C treatment.

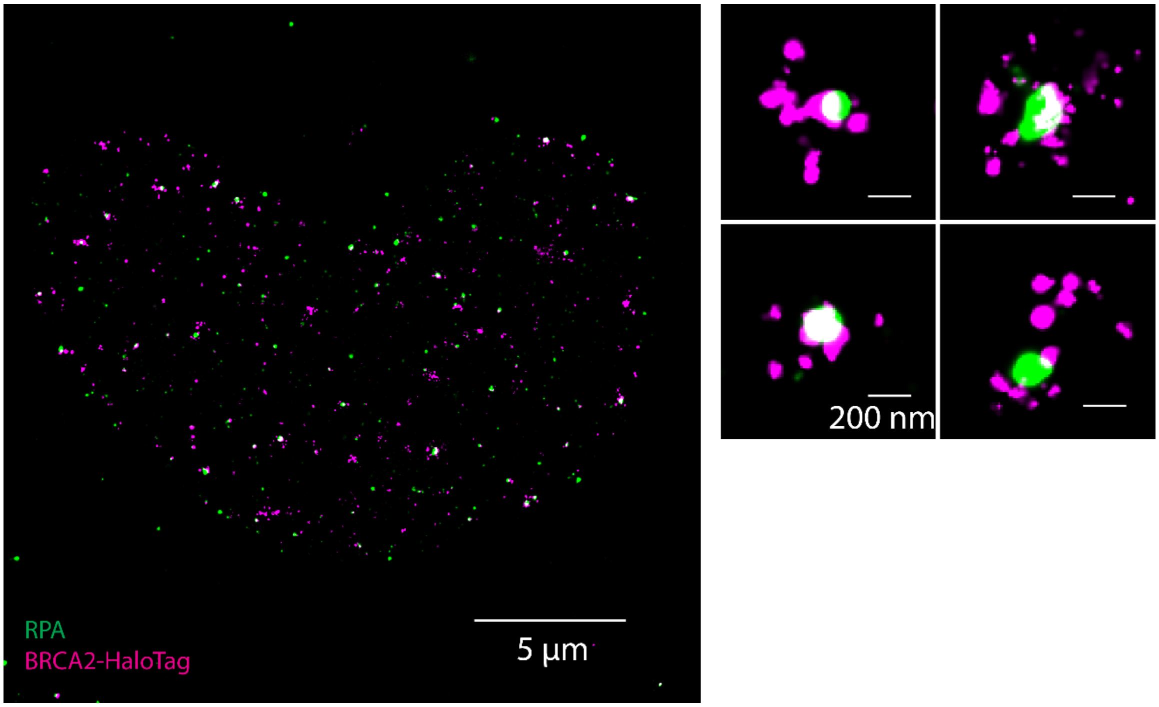


**Fig S7.** dSTORM image of a cell stained for BRCA2-Halo (magenta) with RPA (green), displaying the differential localization of BRCA2 and RPA in repair foci.

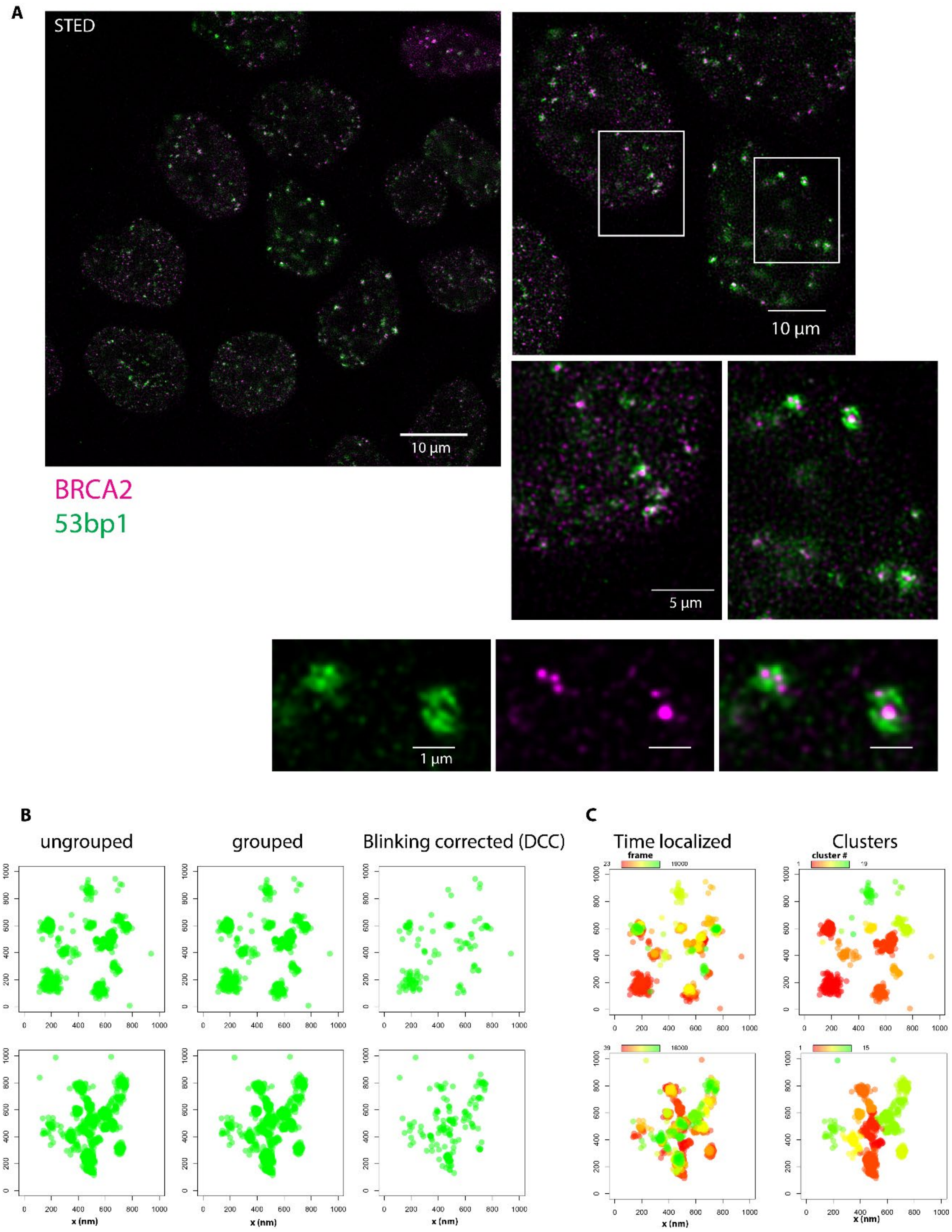


**Fig S8. A** STED image of 53BP1 and BRCA2 in mES cells expressing BRCA2-GFP treated with mitomycin C for 2 hours. Cells are stained with an anti-GFP nanobody with STAR635 (magenta) and anti-53BP1 and a secondary antibody with Alexa594 (green). **B** Example of two BRCA2 repair foci that were imaged with dSTORM from the data in Figure 3. The plots show the localizations of a focus in a 1x1 µm ROI, where dots are individual localizations. The ungrouped plots show the data after localization and drift correction. In grouped localizations are combined which are in subsequent frames and in close proximity. Blinking corrected are the localizations which were obtained using the distance distribution correction (DCC) algorithm to remove blinking artifacts (1). The Time localized plots show the frame in which the grouped localizations were detected in the original dSTORM acquisition. The plot with the clusters shows an example of the clustering as is applied to the dSTORM data in Figure 3 and Figure 4.

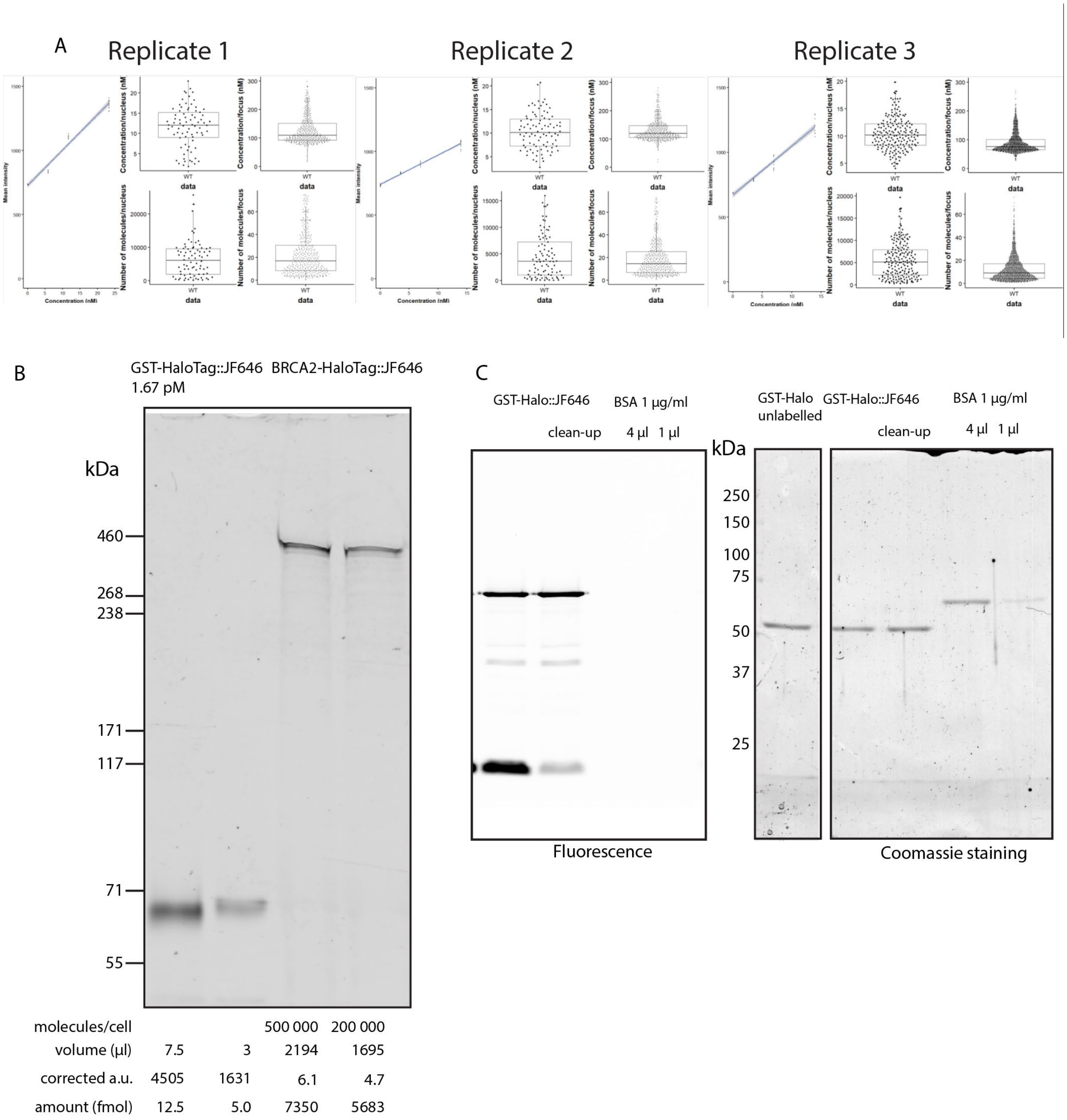


**Figure S9. A** Results of BRCA2-HaloTag quantification in knock-in cell lines treated with mitomycin C. Results of 3 replicates are shown of which the results of replicate 3 are shown in Figure 4. **B** Using a GST-HaloTag protein standard the number of molecules per cell and concentration was determined on a SDS page gel. **C** SDS-PAGE gel showing the results of labelling of the GST-HaloTag reference protein. Detected by fluorescence and by Coomassie staining to validate the concentration of the labeled protein.

**Movie S1.** Example raw data single-molecule movie of BRCA2-Halo in a mitomycin C treated cells with 9 planes projected on the camera

**Movie S2**. Overlay of BRCA2-Halo single-molecule movie and interpolated EGFP-tr53bp1 mask in mitomycin C treated cell

**Movie S3.** Overlay of BRCA2-Halo single-molecule tracks and interpolated EGFP-tr53bp1 mask in mitomycin C treated cell

#### References

1. C. H. Bohrer, *et al.*, A pairwise distance distribution correction (DDC) algorithm to eliminate blinking-caused artifacts in SMLM. *Nat Methods* **18**, 669–677 (2021).
